# Neural mechanisms of perceptive and affective processing of body stimuli in Anorexia nervosa – are there developmental effects?

**DOI:** 10.1101/462952

**Authors:** Stefanie Horndasch, Julie Roesch, Oliver Kratz, Anne Vogel, Hartmut Heinrich, Holmer Graap, Gunther H. Moll, Arnd Dörfler, Clemens Forster

## Abstract

**Objective:** Different components of body image processing seem to be reflected by different neural mechanisms. A core symptom of Anorexia nervosa (AN) is a disturbance of body image with correlates found on a neural level. The present study focusses at the neural processing of visual body stimuli of different weight categories in adolescent and adult AN patients.

**Method:** 33 adolescents aged 12–18 years (15 AN patients, 18 control participants) and 36 adult women (19 AN patients, 17 control participants) underwent functional magnetic resonance imaging (fMRI) while performing a perceptive and an affective body image task involving photographic stimuli of women belonging to different BMI categories.

**Results:** Differential effects on activation depending on the BMI of the women shown in the pictures were found in frontal brain regions, the thalamus, caudate and the fusiform gyrus. Group effects differentiating between AN patients and control participants were seen mainly in the caudate and insula.

**Discussion:** During a perceptive task, diminished activation of regions involved in perceptive and evaluative functions as well as emotional reasoning was seen in AN. During evaluation of the neural processing in an affective task there was a tendency towards activation differences reflecting reduced ability of size estimation and impaired integration of visual and body perception with emotions.

## 1. Introduction

A core symptom of Anorexia nervosa (AN) is body image disturbance which is comprised of a behavioral, perceptual, cognitive and affective component (1-4). Several regions in the human cortex have been investigated as selectively processing the appearance of human bodies in healthy adults, e.g. the “extrastriate body area” in the occipitotemporal cortex (5) or the "fusifom body area" (6). In eating disorders (EDs) abnormal processing of body image, specifically within a neural network for body perception, has been discussed (7) and investigated in several functional Magnetic Resonance Imaging (fMRI) studies. Activation patterns while watching one’s own body (8-13) as opposed to those while looking at others’, mainly female, bodies (9-12, 14-18), have been studied. On top of instructions to simply look at the bodies, specific tasks like weight rating (14, 15, 19) or assessing subjective emotions (11, 14-16, 18) enabled researchers to look at differential aspects of body picture processing.

Differences between AN patients and healthy participants in body image processing, which have been found in the aforementioned studies, can be allocated to the following neural structures:

- In areas which have been described as selectively processing body pictures such as the fusifom body area and extrastriate body area (see above) either enhanced (10) or reduced (17, 18) activity was found in AN patients. However, other studies reported similar levels of activity (9, 16, 20).
- The prefontal cortex (PFC) has been shown as selectively activated when processing others’ bodies in ED patients as well as healthy participants (12, 16, 17, 21), but also to a greater extent in AN patients (9, 10). This is possibly related to the process of mentalizing about oneself. Frontal areas have been considered as being involved in processing self-relevant information (21). Their activity has been linked to body dissatisfaction (19, 22).
- Parietal brain regions have been found to play a role in the processing of body pictures (12, 16, 18, 20, 23). This has especially been seen in AN patients when confronted with one’s own body (23), probably linked to the processing of visospatial information (24). When looking at line drawings of bodies, reduced parietal activity in AN patients has been seen (18).
- Concerning the cingulate cortex, results are more consistently showing reduced activity in AN patients as compared to control participants (16, 20, 21, 25), e.g. when performing a self-“ideal” comparison with pictures of fashion models (16).
- The anterior insula was strongly activated in AN patients when comparing their own body to slim bodies (16) and when viewing distorted thin self portraits and rating their satisfaction (19). By contrast, the posterior insula has been identified as belonging to a subnetwork of regions with decreased connectivity in AN patients (26).
- The ventral striatum, a region which has been linked to reward reactions, shows a different pattern of activation in AN patients and control participants when confronted with women’s bodies of different BMI categories: activation was higher during the processing of underweight stimuli compared with normal weight stimuli in women with AN, and the reverse pattern was observed in healthy females (14).

The latter study was conducted with adult AN patients and later enhanced by an adolescent AN group of which data were analyzed separately. As AN typically has its onset in adolescence (27) and adolescents are highly vulnerable to body image concerns, it is interesting to study the processing of body-related information in adolescent samples in direct comparison to adults. Concerning subjective evaluation of body pictures, age differences were found with adult AN patients showing more aberrant ratings (as compared to healthy participants of the respective age groups) than adolescent AN patients (28). Therefore it is of interest whether neural responses to the pictures used in that evaluation study similarly differ across age groups.

To our knowledge, only two studies differentiated between two aspects of body image processing using a more perceptive and a more affective task (14, 19). Our aim was to examine changes in brain activation when participants are asked to estimate the size and the subjective emotions with respect to different body shapes presented with a standardized picture set.

## 2. Methods

### 2.1. Participants

In total, 69 female participants were included: 33 adolescents (12-18 years) and 36 adults (19-40 years). The patient group consisted of 34 girls and women fulfilling the ICD-10 criteria for anorexia nervosa (ICD-10 F50.0 (29)), the control group of 35 typically developing adolescent girls and healthy adult women in the same age range.

AN patients were recruited at the Department of Child and Adolescent Mental Health and the Department of Psychosomatics and Psychotherapy at the University Clinic Erlangen at the beginning of an inpatient treatment program. Control participants were recruited through advertisements and personal contact. Exclusion criteria for the control group were current or a history of psychiatric disorder and obesity. Further exclusion criteria for all participants were learning disabilities, severe somatic, including neurological disease, a history of head trauma, visual impairments, metallic implants, claustrophobia, other MRI contraindications and psychotropic medication other than selective serotonin reuptake inhibitors or atypical neuroleptics.

All participants completed a German version of the Eating Disorder Inventory-2 (EDI-2), which assesses ED psychopathology and is comprised of 91 items divided into 11 subscales (30). Further, the Physical Appearance Comparison Scale (PACS), a five-item scale that assesses an individual’s tendency to compare their own appearance to the appearance of others (31), and the Sociocultural Attitudes Towards Appearance Questionnaire (SATAQ), designed to assess societal and interpersonal aspects of appearance ideals (32), were applied. Written informed consent was obtained from all participants and from the adolescents’ parents.

The Ethics Committee of the University Hospital of Erlangen gave approval for the study and it was conducted in accordance with the Declaration of Helsinki.

### 2.2. Stimuli

Standardized greyscale photographic stimuli showing women’s bodies in underwear in four positions (front, rear, profile standing and profile sitting) were used (Horndasch et al., 2015). Five pictures from each BMI category were shown: extremely underweight (BMI 13.5–15.0 kg/m^2^), underweight (BMI 17.0–18.0 kg/m^2^), normal weight (BMI 20.0–22.5 kg/m^2^) and overweight (BMI 36.0 – 38.0 kg/m^2^). Examples are presented in Figure 1.

**Figure 1:**
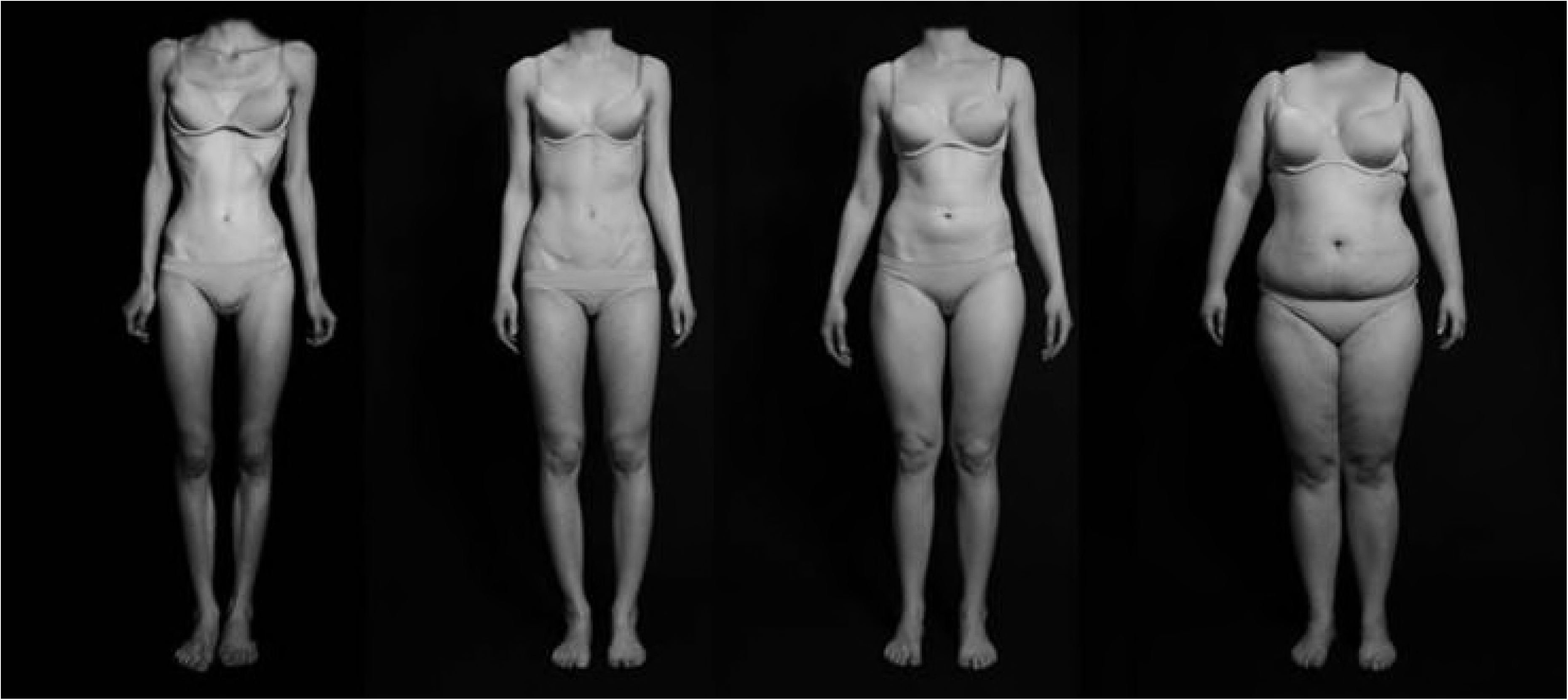
Examples of body stimuli (categories extremely underweight, underweight, normal weight and overweight).

### 2.3. Procedure

Scans were conducted between 4.30 and 6pm. Images were presented via goggles fitted to the head coil (NordicNeuroLab Visual System). Pictures were shown during two tasks in randomized order for 12 sec each with an interstimulus interval of 12 sec. To differentiate between a perceptive and affective component of body image, two different tasks were applied: During the first task (perceptive task), participants had to rate the shape of the body on a 9-point scale ranging from “very underweight” to “very overweight”. During the second task (affective task), participants were asked to process the stimuli in a self-referential way and imagine themselves having the body shape of the women displayed in the picture. They were asked to rate their satisfaction with this imaginary body shape on another 9-point scale ranging from “very unhappy” to “very happy”. Ratings were performed between picture presentation and the interstimulus interval and had a fixed duration of 6 sec. The total duration of each task was therefore 10 minutes. Two participants performed only the perceptive, but not the affective task due to technical problems.

### 2.4. Image Acquisition

All data was acquired using a 12-channel head coil on a 3T high-field scanner (Magnetom Tim Trio, Siemens Healthcare AG, Erlangen, Germany). Anatomical images were acquired using a T1-weighted 3D MPRAGE sequence (1 mm isotropic resolution, TR/TE/FA = 1900/2.25/9, Field of view 25.6 cm × 25.6 cm × 25.6 cm). Functional MRI data (depicting Blood Oxygen Level Dependent (BOLD) contrast) were acquired using a multi-slice 2D EPI sequence (1.5 mm × 1.5 mm in plane resolution, 3 mm slice thickness, 0,75 mm interslice gap, TR/TE/FA = 3000/30/90, FOV = 192mm, 128 × 128 matrix). 400 T2-weighed whole brain volumes were acquired [for an investigation of processing of food and emotional stimuli with this method see (33)].

### 2.5. Data Analysis and Statistics

Behavioral data (subjective valence ratings) were analyzed by using an analysis of variance (ANOVA) and post hoc two-tailed t tests using Bonferroni-Holm correction to control for multiple comparisons.

Imaging data analysis, registration, visualization and statistical analyses were performed with the BrainVoyager QX software package (Brain Innovation B.V., Maastricht; The Netherlands). Pre-processing of the data included three-dimensional motion correction, temporal Gaussian smoothing of 4 sec, spatial Gaussian smoothing of 4 mm, linear detrending and temporal high pass filtering using 0.01 Hz. To better assign the brain activities to anatomical locations, the individual brains were transformed into standard stereotactic space (Talairach and Tournoux, 1988) and the functional data were co-registered and superimposed with the anatomical data (3-dimensional MPRAGE data set). Talairach coordinates were used to confirm anatomical regions of activation using Talairach Daemon (Research Imaging Center, University of Texas Health Center, San Antonio, TX, USA). Analysis was done with a general linear model with subjects as a random effect. A boxcar-like predictor was used which described the stimulus category (extremely underweight, underweight, normal weight or overweight). The predictor function was convolved with a haemodynamic response function to consider delays and graded response times of the BOLD signal to the stimuli. For this a 2γ-haemodynamic response function was used (time to response to peak: 5 s, time to undershoot peak: 15 s). The effects of the predictors on cortical activation were analyzed for AN patients and control participants.

The analysis was restricted to regions of interest (ROIs) which had previously been described as being involved in the processing of body stimuli (see Table 1).

**Table 1.**
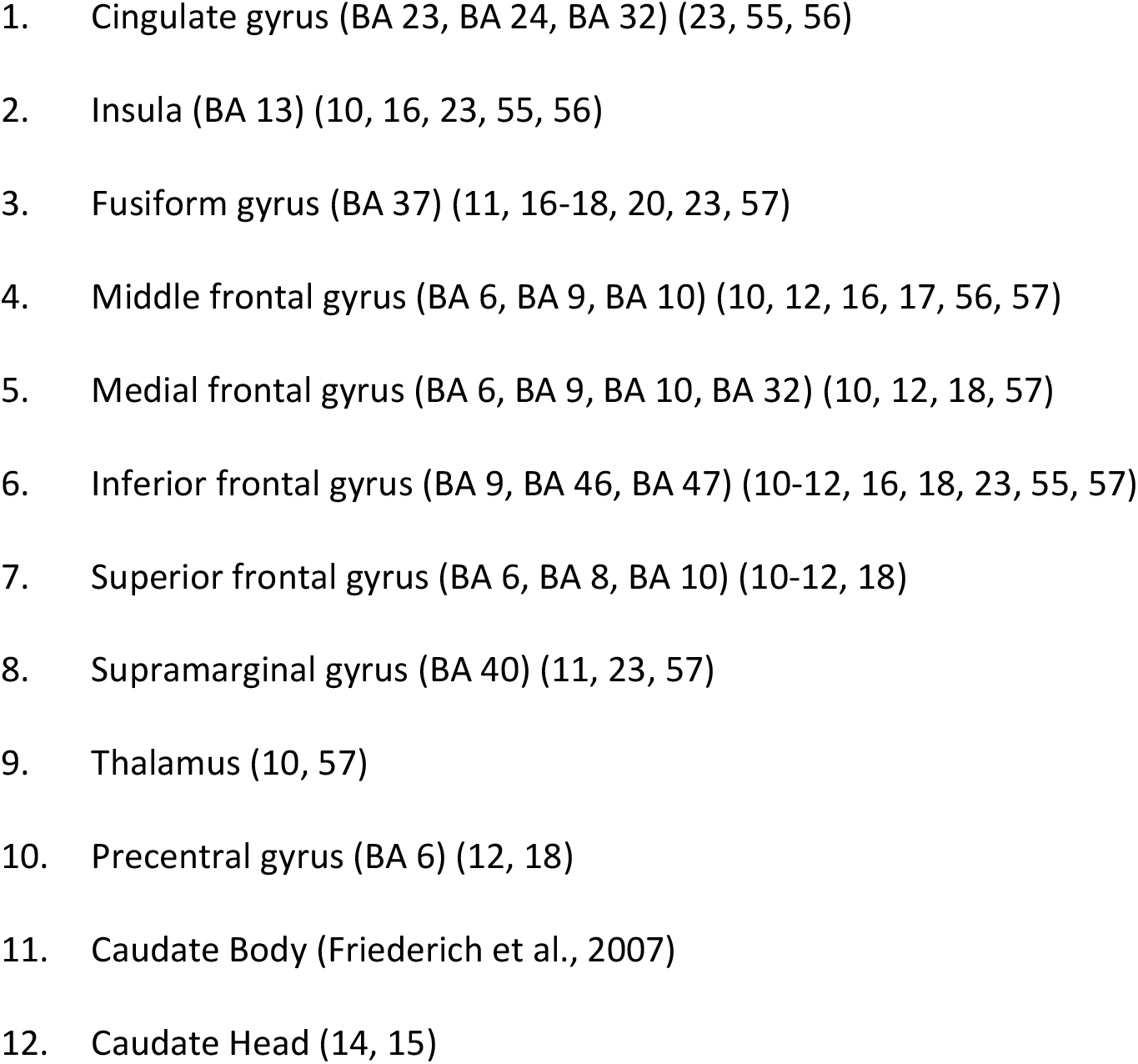
Regions of Interest. Selected Regions of Interest based on previous studies on processing of body stimuli in EDs and in the developing brain.

A cortical brain site was regarded as activated only if the significant change of the BOLD signal covered a minimum cluster size of 150 mm^3^ (see e.g. (34, 35)). The level of significance for the detection of activated cortical areas was determined by Bonferroni correction for the number of regions of interest (ROI) analyzed. 46 ROI were defined which yielded a level of significance of *α*’=0011. Only clusters which belonged to one of the ROIs defined were taken for the ongoing analysis.

The strength of the activation within subjects’ ROIs was expressed by the individual t values which were obtained by a second level analysis with the subject as a factor. Analyses were performed at the group level (multi-study t tests) for adult and adolescent participants separately.

In order to clarify effects of picture category on brain activation patterns rather than merely group differences of processing of the single categories, individual β values were extracted for those regions in which group differences (patients vs. control participants) were evident (see Table S1 for regions with group differences; a detailed description of those group statistics would be beyond the scope of the present paper). These β values were used as a measure of the strength of the individual BOLD changes within the respective ROI. They were then subjected to a repeated measures ANOVA with “BMI category” of the stimuli (extremely underweight, underweight, normal weight and overweight) as within-subjects variable and “participant group” (2, “AN patients” and “control participants”) and “age group” (adolescents vs. adults) as between-subject factors. Greenhouse-Geisser correction was used when appropriate. For brain regions with significant “participant group” or “age group” interactions, group comparisons via t tests were conducted using Bonferroni-Holm correction to control for multiple comparisons. For some regions that we considered to be of importance for the study the effects were only close to significance. These cases were treated and discussed as tendencies. For regions with significant main effects, planned comparisons were performed in order to determine activation differences between groups for the single picture categories, Bonferroni-Holm correction was also applied. The ANOVA and post hoc tests were carried out using SPSS 21.0 (IBM Corp.).

## 3. Results

### 3.1. Participant characteristics

There were no group differences between AN patients and control participants regarding age for adolescents as well as for adults. BMI (adults) and BMI age percentile (adolescents) were significantly lower for AN patients. ED symptoms assessed by the EDI-2 total score and drive for thinness subscale, depressive symptom scores assessed by the BDI and the tendency to internalize appearance ideals assessed by the SATAQ were higher in AN patients with larger group differences in adults than in adolescents. For adults, but not for adolescents, group differences were seen in the appearance comparison behaviour assessed by the PACS and for body dissatisfaction (see Table 2). Three adolescent and nine adult AN patients were taking antidepressant medication and one adolescent and three adult patients neuroleptic medication at the time of scanning. Comorbidities in the AN sample were depressive disorder (13 adolescents and 13 adults), obsessive-compulsive disorder (one adolescent, one adult), anxiety disorder (four adolescents and two adults) and post-traumatic stress disorder (two adults).

**Table 2.** Participant characteristics. Mean, standard deviation and statistical p-value of the difference between both groups are reported.

| Age group | Adolescents (n = 33) |  |  | Adults (n = 36) |  |  |
| --- | --- | --- | --- | --- | --- | --- |
| Participant group | AN (n=15) | Control participants (n=18) | p | AN (n=19) | Control participants (n=17) | p |
| Age (years) | 15.72 (1.93) | 16.45 (1.35) | n.s. | 26.27 (6.71) | 27.03 (5.46) | n.s. |
| BMI (kg/m<sup>2</sup>) |  |  |  | 16.06 (2.04) | 21.40 (1.40) | < .001 |
| BMI age percentile | 5.33 (11.67) | 56.33 (25.48) | <.001 |  |  |  |
| SATAQ | 53.69 (17.31) | 40.71 (11.96) | < .05 | 54.24 (13.37) | 39.46 (11.04) | <.01 |
| PACS | 15.85 (4.76) | 14.29 (2.95) | n.s. | 13.00 (3.89) | 17.65 (3.20) | < .001 |
| BDI | 20.77 (12.41) | 10.59 (9.61) | < .05 | 29.50 (8.44) | 5.87 (7.95) | < .001 |
| EDI-2 total score | 283.57 (57.08) | 248.47 (37.59) | < .05 | 337.47 (49.04) | 208.47 (54.4) | < .001 |
| EDI-2 drive for thinness | 18.82 (7.17) | 28.4 (9.91) | < .01 | 32.71 (7.83) | 15.69 (4.44) | < .001 |
| EDI-2 body dissatisfaction | 31.13 (12.1) | 29.65 (4.92) | n.s. | 43.41 (6.5) | 29.44 (9.22) | < .001 |

### 3.2. Ratings

#### 3.2.1. Perceptive task

Regarding weight ratings of women’s bodies of the four different categories, the ANOVA revealed the following significant effects: A “BMI category” main effect (F(3,189)=1238.8, p<.001, part. η^2^=.952) indicates that weight ratings increased with increasing BMI of the women shown in the pictures. Further we found a main effect of “participant group” (F(1,63)=9.5, p<0.01, part. η^2^=.132) with higher weight ratings by AN patients. A significant “age group”*”participant group” interaction (F(1,63)=4.7, p<0.05, part. η^2^=.069) indicates that the effect of AN patients estimating the women’s BMI as being higher than controls is stronger in adults than in adolescents.

Group comparisons via t tests showed that the weight of extremely underweight and normal weight women in the pictures were rated higher by adult AN patients than by healthy women. In adolescents, no significant group differences for the single categories emerged after Bonferroni-Holm correction (see Figure 2).

**Figure 2:**
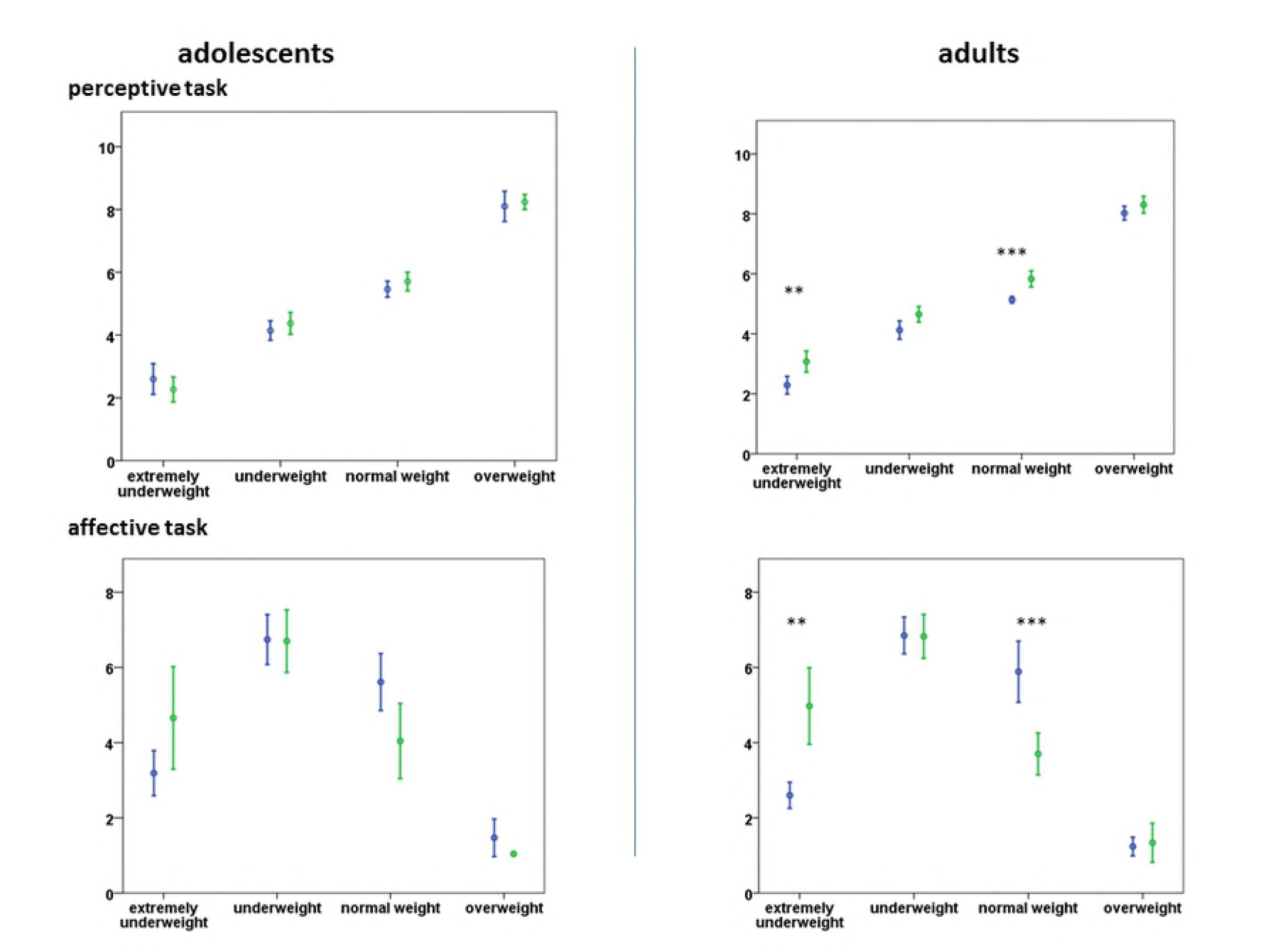
Subjective ratings of adolescent and adult patients in the perceptive and affective task (means ± 2 standard deviations are shown). Significant group differences are indicated: **=p<.01, ***=p<.001 after Bonferroni-Holm correction.

#### 3.2.2. Affective task

In affective ratings a “BMI category” main effect (F(3,180)=161, p<0.001, part. η^2^=.729) with highest ratings for underweight, followed by normal weight, extremely underweight and overweight body shapes was seen. A significant “BMI category”*“participant group” interaction (F(3,180)=19.9, p<0.001, part. η^2^=.249) indicated higher satisfaction ratings in the control group for pictures of normal weight women compared to pictures of the extreme underweight category, but not in AN patients.

Direct group comparisons within the age groups revealed higher satisfaction ratings for extremely underweight bodies in adult AN patients and lower ratings for normal weight bodies. Adolescent AN patients rated their satisfaction for all weight categories similarly to control participants (see Figure 2).

### 3.3. Activation patterns

Beta values of activated clusters were subjected to separate ANOVAs for the perceptive and the affective task respectively. Significant effects in brain regions previously seen as involved in body image processing were found for “BMI category” of the women in the pictures, “participant group”, “age group” and “participant group” and “BMI category” interaction (see Table 3 for significant effects). In the perceptive and affective task effects of differential activation occurred partly in similar regions (e.g. frontal areas, fusiform gyrus, Thalamus), partly in different areas (e.g. supramarginal gyrus, insula, caudate body and head).

**Table 3.** Significant main effects and interactions (ANOVA with the factors “BMI category”, “participant group” and “age group”) for the perceptive and affective task after Bonferroni correction (p’<0.0011). BA: Brodman area, TC: Talairach coordinates, F: F statistic, *:p<.05, **:p<.01, ***:p<.001, °:<.1 after Bonferroni correction.

Perceptive Task
|  |  | BA | laterality | TC |  |  | F |
| --- | --- | --- | --- | --- | --- | --- | --- |
| Main „BMI category“ effect | Thalamus, ventral lateral nucl. |  | R | 12 | -14 | 9 | 12.99*** |
|  | Middle frontal gyrus | 9 | R | 46 | 31 | 34 | 19.22*** |
|  | Middle frontal gyrus | 10 | R | 39 | 51 | 10 | 11.90*** |
|  | Medial frontal gyrus | 32 | R | 9 | 11 | 45 | 8.81** |
|  | Inferior frontal gyrus | 9 | R | 50 | 0 | 24 | 8.89** |
|  | Inferior frontal gyrus | 46 | L | -43 | 7 | 30 | 6.98** |
|  | Inferior frontal gyrus | 47 | R | 50 | 17 | -7 | 8.44** |
|  | Fusiform gyrus | 37 | L | -39 | -48 | -9 | 15.66*** |
|  | Fusiform gyrus | 37 | R | 45 | -56 | -7 | 39.41*** |
| Main „participant group“ effect | Caudate body | 23 | L | -11 | -20 | 26 | 12.77* |
|  | Caudate body |  | R | 9 | 1 | 12 | 19.24** |
| “Participant group”*“BMI category” interaction | Superior frontal gyrus orbital | 10 | L | -31 | 58 | -2 | 10.08*** |
|  | Insula anterior | 13 | L | -34 | 5 | 14 | 5.85* |

**Affective Task**
|  |  | BA | laterality | TC |  |  | F |
| --- | --- | --- | --- | --- | --- | --- | --- |
| **Main „BMI category“ effect** | Thalamus, ventral lateral nucl. |  | R | 12 | -14 | 9 | 7.95** |
|  | Inferior frontal gyrus | 9 | R | 50 | 0 | 24 | 15.69*** |
|  | Inferior frontal gyrus | 46 | L | -43 | 7 | 30 | 10.22*** |
|  | Inferior frontal gyrus rostral | 47 | R | 22 | 35 | -3 | 16.52*** |
|  | Inferior frontal gyrus | 47 | L | -39 | 25 | 0 | 6.33* |
|  | Middle frontal gyrus | 9 | R | 27 | 38 | 25 | 36.39*** |
|  | Middle frontal gyrus | 10 | R | 39 | 51 | 10 | 22.95*** |
|  | Medial frontal gyrus | 32 | R | 9 | 11 | 45 | 14.08*** |
|  | Fusiform gyrus | 37 | L | -39 | -48 | -9 | 13.50*** |
|  | Fusiform gyrus | 37 | R | 45 | -56 | -7 | 52.93*** |
|  | Supramarginal gyrus | 40 | R | 61 | -43 | 29 | 10.20*** |
|  | Caudate head |  | L | -11 | 11 | 1 | 5.96* |
| **Main „age group“ effect** | Caudate Body | 23 | L | -11 | -20 | 26 | 11.71° |
| **“Participant group”\*“BMI category” interaction** | Insula posterior | 13 | R | -42 | -15 | 19 | 5.65° |

#### 3.3.1. BMI sensitive brain regions

Concerning the “BMI category” main effect, decreasing activity with increasing BMI of the women in the pictures was seen for most regions (thalamus, middle, medial and inferior frontal gyrus, fusiform gyrus). Exceptions were the supramarginal gyrus and the caudate head in the affective task. The former showed stronger activity with growing BMI of the women in the pictures. The caudate head showed decreased activation for the category normal weight and similar activation with beta values close to zero for categories extremely underweight, underweight and overweight.

#### 3.3.2. Group sensitive brain regions

The planned comparison analysis of the main “participant group” effect in the perceptive task revealed group differences between AN patients and control participants in the left caudate body for extremely underweight (t(66)=3.15, p<.01) and normal weight (t(66)=2.68, p<.05) body pictures and in the right caudate body also for extremely underweight (t(66)=3.41, p<.01) and normal weight (t(66)=3.01, p<.05) bodies with a decreased BOLD signal in AN patients and positive values for control participants. In the affective task a tendency close to significance towards an “age group” main effect (p<.06) was found in the left caudate body with higher activity in adolescents than in adults for extremely underweight (t(64)=3.26, p<.01) and underweight (t(64)=2.61, p<.05) pictures.

A “participant group” * “BMI category” interaction in the perceptive task was shown for the superior frontal (orbitofrontal) gyrus as well as the left anterior insula. In the superior frontal gyrus a significant group difference with an increase of the BOLD signal in controls and a decrease in patients was shown for underweight (t(66)=3.56, p<.01) body pictures and a trend towards significance for extremely underweight (t(66)=2.95, p<.1) pictures. In the anterior insula a significant effect in the same direction was evident for extremely underweight pictures (t(66)=4.11, p<.001; see Figure 3). In the affective task, a tendency towards a “participant group” * “BMI category” interaction for the right posterior insula represented significantly higher (and highest of all categories) BOLD signal for extremely underweight (t(64)=4.16, p<.001) pictures in control participants vs. signal decrease for this category and increasing activity with BMI of the bodies in the pictures for AN patients (see Table 3 for ANOVA main effects and interactions).

**Figure 3:**
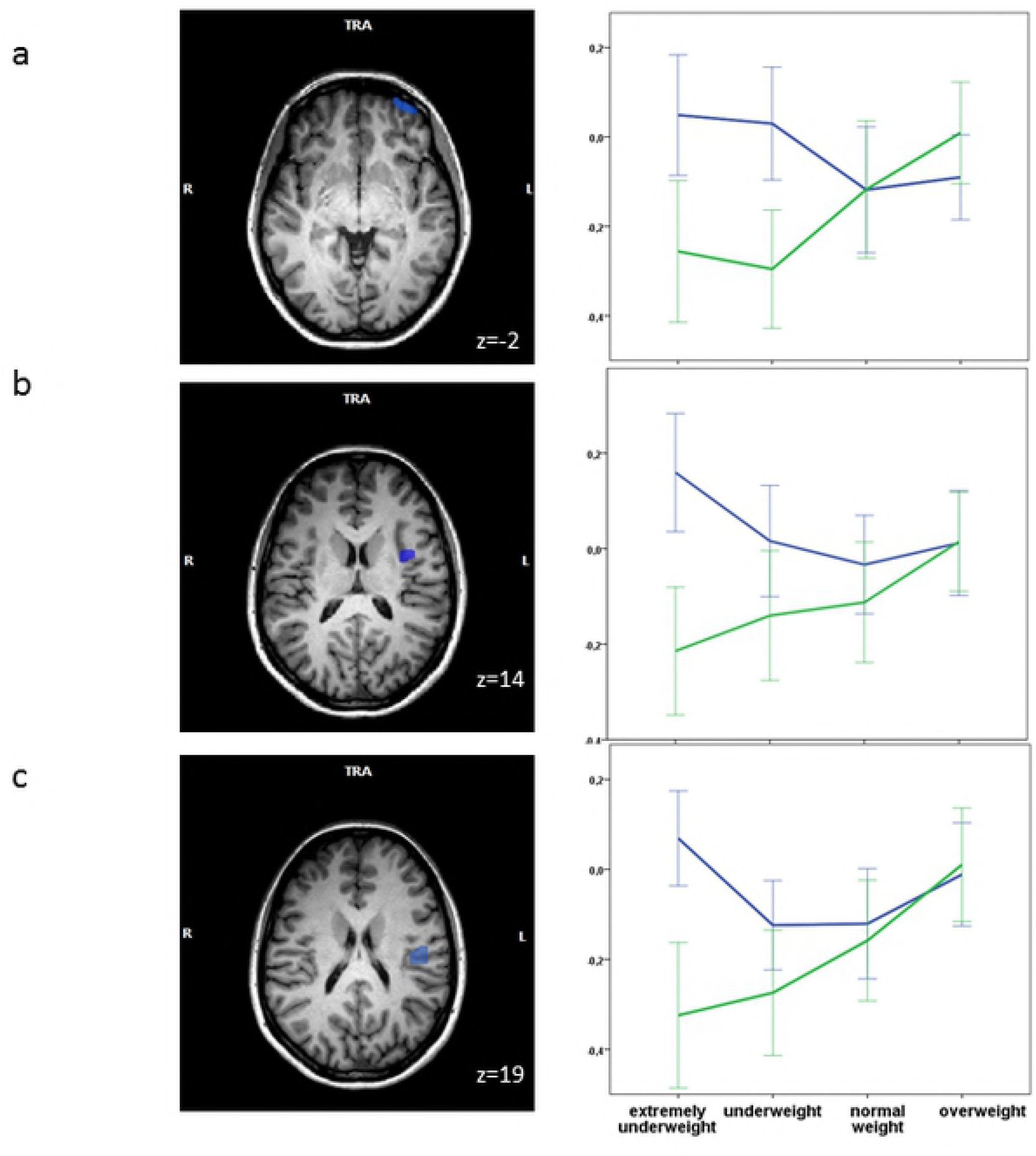
Activation patterns (beta values) in AN patients (green) and control participants (blue) across picture categories (mean ± 2 standard errors).

## 4. Discussion

### 4.1. Subjective Ratings

Women’s bodies were generally rated as having more weight by AN patients than by females not suffering from an eating disorder. However, this effect seems to be mainly based on group differences in adult participants, whereas direct comparisons between adolescent patients and control participants revealed no differences. When asked about their satisfaction with a body of a specific weight category, AN patients showed a bias towards assessing extremely underweight bodies as more desirable (adult patients only) and normal weight bodies as less desirable (adolescent and adult patients) than healthy participants. These effects were similarly shown in a larger patient sample (reported in Horndasch et al., 2015) and – with regard to the latter (satisfaction), but not the former (weight estimation) effect – in a previous study with adult participants looking at pictures of bodies of three different weight categories (14).

### 4.2. Brain activation patterns

#### 4.2.1. BMI sensitive brain regions

In a number of brain regions there were differential effects on activation depending on the BMI of the women shown in the pictures. A region discussed in numerous previous studies is the fusiform gyrus. The BOLD signal, interpreted as activation of this region, was strongest when viewing strongly underweight women with a linear decrease over BMI categories and lowest activation for overweight body stimuli in the perceptive as well as the affective task of our study. Similar regions within the fusiform gyrus were identified by Peelen et al. (2005) to be selective for the visual processing of human bodies (vs. tools) in healthy participants. However, in our results, no group differences like in other studies (7) were found.

Another region discriminating between different weight categories of bodies in both tasks in a similar manner was the right thalamus. In healthy as well as AN participants, significant thalamus activation was seen before when viewing non-self body images (Sachdev et al., 2008) and activity of the left thalamus was linked to word body stimuli in AN patients (Miyake et al., 2010). Specific thalamic reactions towards body cues are therefore a known phenomenon, but previous results didn’t differentiate between weight sub-categories.

Frontal regions also showed a pattern of decreasing activity from extremely underweight to overweight body pictures. Regions belonging to the prefrontal cortex (BA 46, 47) exhibited this pattern in both tasks, but inferior frontal regions (BA 9) more specifically in the affective task. Prefrontal regions have been shown to be activated to a greater extent in AN patients while watching others’ bodies or reading body words (9, 36), but also in control participants (Sachdev et al., 2008). Inferior frontal regions have even more frequently been linked to body image tasks. Looking at bodies vs. chairs caused increased inferior frontal activation in healthy participants (Suchan et al., 2012). Regions of BA 9 have been activated in response to body images, but less to images of interior design (Friederich et al., 2007) and in body-shape self-comparison in AN patients (Friederich et al., 2010) as well as in controls (Sachdev et al., 2008, Wagner et al., 2003) and have been seen in the context of self-reflection and self-judgement (21). Medial frontal regions have been linked to cognitive aspects of emotional processing, paying attention to emotions and fear processing (37, 38). These mechanisms have possibly occurred in all participants in our study when looking at strongly underweight other bodies which are not as common as and probably inducing more (negative) emotions than normal weight or overweight bodies. Generally, the pattern of increased activation of different regions for strongly underweight and, to a lesser extent, underweight bodies could be seen as increased alertness and attention in response to these unusual body shapes. Similarly, increased central nervous responses in the form of enhanced event related potential amplitudes (“Late Positive Potential”) towards strongly underweight bodies were seen in an EEG study using the same stimulus set in an overlapping sample, but occured in AN patients only (39). In contrast to the MRI activation patterns described here, those increased reactions observed at electrodes over central brain areas seem to be more specific to patients, probably reflecting more motivational processes. However, as a number of brain regions are assumed to be generators of the Late Positive Potential, activation patterns in specific brain regions cannot be directly linked to enhanced amplitudes.

The caudate head (ventral striatum) also showed a specific pattern regarding the different weight categories in the affective task. In the studies of Fladung et al. (2010, 2013) there was also a greater effect for this region when participants fulfilled an affective than a mere perceptive task with enhanced activation for underweight in comparison to normal weight body pictures in AN patients. In our study this reward-related brain structure did not show greatest activation for extremely underweight bodies in AN patients and no group effect was found. Maybe the different stimulus types used in the studies might play a role and computer generated underweight pictures – as used in the Fladung et al. studies – might have looked less realistic and therefore more “rewarding” and less deterring for AN participants.

The supramarginal gyrus as the only exception in weight category-sensitive brain regions revealed by our study displayed growing activity with growing BMI of the women in the pictures with a clear peak for overweight bodies. This region in the parietal cortex is said to be part of the somatosensory association cortex and involved in identifying bodily postures (40) and proprioception (41). Lower activation in the inferior parietal cortex in AN patients than in controls was reported in different tasks (10, 42) and connected to disturbances in visuo-spatial processing (23), while in our study no group differences could be found in this region.

#### 4.2.2. Group sensitive brain regions

For the perceptive task, the left and right caudate body differentiated healthy women from women with acute anorexia nervosa. The caudate has been highlighted as being important in food cue processing with distinctive patterns – mainly in the direction of hyper-activation – in AN (43, 44). However, the caudate body has also been linked to perceptual processes with input from other perceptive regions (45) and is also involved in body perception and judgement in healthy participants (46). In a study on self-comparison with model pictures in AN patients, a positive relationship between subjective anxiety in response to body images and activation in the caudate body was seen (21). However, in our body size estimation task in which participants were probably less emotionally involved, control participants showed higher caudate activity than AN patients. This finding could – together with the fact of regional decreases in gray and white matter which have been shown for this area in AN patients (47) – hint towards reduced perceptive and evaluative functions regarding human bodies in AN.

Another group specific pattern in the perceptive task was seen in the orbitofrontal gyrus (BA 10), with decreased activity for the underweight BMI category for AN patients. Activity in BA 10 correlates positively with body dissatisfaction in AN (22). So possibly underweight pictures cause, relative to other categories, less dissatisfaction in AN. The region has also been linked to emotional reasoning (48), which could be impaired when estimating strongly underweight and underweight bodies in AN.

For the anterior insula a similar distribution has been found, with diminished activity in AN patients, especially for extremely underweight bodies, vs. highest activity for those emaciated bodies in control participants. Again, this region was mainly found to be functionally affected in AN patients in satisfaction rating tasks related to slim images (16, 19), but is showing a different effect in our size estimation task. The anterior insula may represent strive for thinness and motivational tendencies, which does not seem to be consistent with increased activation for higher BMIs in our study, but is also part of the fear and emotional network (16). Therefore activation decrease for extremely underweight bodies in patients might point towards an impaired integration of visual and body perception with emotions, especially fear, which might play a role in control participants, but not in AN patients. The insula is connected with many structures for emotional processing and bodily experience, among those frontal regions and the striatum which show reduced activity in our task as well (see above). Dysfunction within the fronto-insula-limbic-striatal circuit has been discussed in the context of failure to regulate inputs from higher level cognitive representations about the importance of weight and shape (49).

By contrast, in the affective task, the posterior insula showed a tendency towards a group specific effect with highest activation for “extreme” BMI categories (extremely underweight, overweight) in control participants vs. increasing activity with BMI of the bodies in the pictures for AN patients and a significant difference for extremely underweight shapes. In spite of having received less attention than the anterior section in the previous literature, the posterior section of the insula has also been discussed in the context of eating disorders. It has been identified as part of a subnetwork of connections with decreased connectivity in AN and discussed as a central “hub” within this network (26). Specifically the posterior insula has been associated with sensorimotor processing with functional connections to motor and somatosensory cortical areas and may thus be involved in the processing of own bodily signals such as body size (Ehrlich, Lord et al. 2015). Larger involvement for extreme body shapes in healthy participants might reflect salience of those shapes. On the contrary, reduced activity in AN patients might be due to an impairment of size estimation, especially when it comes to dangerously underweight shapes similar to the patients’ own.

#### 4.2.3. Developmental perspective

A strong tendency towards a difference in activation patterns between adults and adolescents was found for the affective task in the caudate body. Here generally higher activation, specifically with significance for (extremely) underweight pictures, was seen in adolescents as compared to adults. To date, few studies on body image processing have been conducted with adolescent participants. In our EEG study, we saw differences between adolescents and adults in the form of increased responses towards extremely overweight bodies in adolescents (39). Regarding brain activation patterns, however, no differences regarding overweight bodies were found in this study. As discussed above, the caudate body is involved in perceptual processes, but also connected to anxiety in a self-comparison condition (21). During adolescence, the striatum is known to undergo maturational changes in accordance with changes in emotionality, social interactions and risk-taking (50). The region receives dopaminergic input from the substantia nigra and ventral tegmental area, forming an important part of the reward network. Upon the receipt of reward, the striatum response is stronger in adolescents compared to children and adults (51), and the region is also involved in adolescent social reinforcement learning (52). In adolescents, there is an activity increase in the striatum associated with the expectation to be liked by a peer (53). Therefore the connection to peer evaluation and different emotions related to such a process, which is often linked to ideas about one’s own appearance and body shape during adolescence, could lead to an increased caudate activation in adolescents compared to adults. This could be especially relevant when looking at underweight shapes, which are more likely to be targets for social comparison and evaluation processes.

## 5. Conclusion

In a task using representations of bodies belonging to different BMI categories, several brain regions differentiated well between the distinct categories and therefore appear to be involved in body size assessment. During size perception of the bodies, there was underactivation of regions involved in perceptive and evaluative functions and decreased anterior insula activation in AN, possibly representing an impaired integration of visual and body perception with emotions, especially fear. During an emotional task, a tendency towards reduced posterior insula activation in AN might represent impairment of size estimation for extremely underweight shapes and disturbances in regulatory networks. A trend towards differences between adolescents and adults might show a larger importance of evaluative and comparative processes in young participants. For future research, it would be interesting to compare different tasks beyond merely perceptive and affective components, e.g. using elements of reframing or acceptance therapy to manipulate and evaluate participants’ attitude towards their body. It would also be interesting to acquire longitudinal data in the course of treatment and, in order to further clarify the processes involved, to assess neural activation patterns in the context of e.g. dopamine depletion, as dopamine has been seen to influence reward reactions in AN (54).

## Supplementary material

**S1 Table.** Results of the contrast analysis of activation differences between adult and adolescent AN patients and control participants (T values with significant group differences in p values – p’<0.0011 – in bold print). S1_1: Perceptive Task, adults

| Contrasts |  |  |  |  |  | T-Values |  |  |  |
| --- | --- | --- | --- | --- | --- | --- | --- | --- | --- |
| Region of Interest |  |  | x | y | z | extreme underweight | underweight | normal weight | overweight |
| Anterior cingulate | 24 | R | 3 | 22 | 10 | -1,757 | -0,487 | -0,221 | -0,936 |
| Cingulate gyrus | 32 | L | 3 | 48 | 2 | **4,260** | 3,227 | -0,287 | **4,078** |
| Insula posterior | 13 | R | 46 | -17 | 18 | -2,791 | **-3,684** | -1,296 | 1,336 |
| Insula anterior | 13 | L | -34 | 5 | 14 | -2,214 | -1,78 | 0,253 | -0,247 |
| Insula posterior | 13 | L | -42 | -15 | 19 | **-3,67** | -1,5 | 0,133 | -0,713 |
| Caudate head |  | L | -11 | 11 | 1 | -1,562 | -1,725 | -1,613 | -1,711 |
| Middle frontal gyrus | 9 | L | 39 | 38 | 30 | 1,601 | 1,625 | **4,137** | 0,146 |
| Inferior frontal gyrus | 9 | R | 50 | 0 | 24 | 2,746 | 1,829 | 1,796 | 2,476 |
| Middle frontal gyrus | 9 | R | 46 | 31 | 34 | 2,297 | 0,068 | 3,101 | 0,266 |
| Superior frontal gyrus orbital | 10 | L | -31 | 58 | -2 | -1,85 | -1,364 | **3,205** | 2,63 |
| Medial frontal gyrus | 10 | L | -10 | 65 | 0 | 2,659 | 2,006 | **5,06** | 1,889 |
| Middle frontal gyrus | 10 | R | 39 | 51 | 10 | 0,328 | -0,759 | 2,072 | -0,126 |
| Medial frontal gyrus | 32 | R | 9 | 11 | 45 | 2,263 | 1,338 | 2,7 | 1,242 |
| Medial frontal gyrus | 6 | L | -5 | 2 | 54 | **4,673** | 3,704 | **4,586** | 1,786 |
| Superior frontal gyrus | 6 | R | 1 | -3 | 72 | 0,165 | -0,876 | 0,099 | 2,422 |
| Precentral gyrus | 6 | L | -39 | -9 | 66 | 2,103 | 1,759 | **3,86** | 0,109 |
| Middle frontal gyrus | 6 | R | 34 | 6 | 45 | **4,673** | **3,704** | **4,586** | 1,786 |

S1\_2: Perceptive Task, adolescents
| Contrasts |  |  |  |  |  | T-Values |  |  |  |
| --- | --- | --- | --- | --- | --- | --- | --- | --- | --- |
| Region of Interest |  |  | x | y | z | extreme underweight | underweight | normal weight | overweight |
| Caudate body |  | L | -11 | -20 | 26 | **-6,142** | **-3,363** | **-3,903** | -2,19 |
| Posterior cingulate | 23 | R | 8 | -25 | 20 | **-3,825** | **-4,091** | **-3,498** | -0,359 |
| Cingulate gyrus | 24 | R | 8 | 6 | 37 | **-5,748** | -0,797 | -1,486 | 0,521 |
| Anterior cingulate gyrus | 32 | L | 3 | 48 | 2 | -1,194 | 0,358 | -3,193 | 0,224 |
| Insula posterior | 13 | R | 46 | -17 | 18 | **-5,708** | -2,764 | **-3,627** | -1,529 |
| Insula anterior | 13 | L | -34 | 5 | 14 | **-7,275** | -1,912 | -2,223 | 0,276 |
| Insula anterior | 13 | R | 39 | 6 | 14 | **-5,405** | -1,92 | -1,692 | -0,422 |
| Insula posterior | 13 | L | -42 | -15 | 19 | -3,036 | -0,203 | -1,661 | 1,091 |
| Fusiform gyrus | 20 | R | 46 | -29 | -25 | -2,528 | -2,27 | -2,207 | -1,64 |
| Caudate body |  | R | 14 | -17 | 25 | **-6,136** | -1,157 | **-3,884** | -2,997 |
| Middle frontal gyrus | 9 | L | 39 | 38 | 30 | **-3,567** | 0,422 | -1,027 | 0,444 |
| Inferior frontal gyrus | 9 | R | 50 | 0 | 24 | 1,604 | **5,643** | 2,003 | -0,113 |
| Middle frontal gyrus | 9 | R | 46 | 31 | 34 | 0,735 | 3,159 | 1,893 | -0,621 |
| Superior frontal gyrus orbital | 10 | L | -31 | 58 | -2 | **-5,79** | **-6,483** | -3,12 | -0,449 |
| Superior frontal gyrus | 10 | R | 32 | 56 | -3 | **-3,406** | **-4,691** | -2,012 | -1,383 |
| Middle frontal gyrus | 10 | R | 39 | 51 | 10 | -2,357 | -0,452 | -1,361 | -1,446 |
| Medial frontal gyrus | 32 | R | 9 | 11 | 45 | **-5,154** | -0,798 | -1,191 | -1,193 |
| Supramarginal gyrus | 40 | R | 61 | -43 | 29 | -0,358 | **3,59** | 1,959 | 1,171 |
| Inferior frontal gyrus | 9 | L | -43 | 7 | 30 | -1,78 | 1,995 | -0,123 | 0,904 |
| Inferior frontal gyrus | 47 | L | -39 | 25 | 0 | **-4,446** | -0,655 | -0,682 | -0,761 |
| Inferior frontal gyrus rostral | 47 | R | 22 | 35 | -3 | 1,304 | 2,017 | 3,065 | -0,529 |
| Thalamus ventral lateral nucleus |  | R | 12 | -14 | 9 | -4,273 | 0,469 | -2,642 | -0,046 |
| Medial frontal gyrus | 6 | L | -5 | 2 | 54 | **-5,912** | 0,818 | -0,226 | 0,016 |
| Superior frontal gyrus | 8 | L | -18 | 20 | 47 | -1,516 | -1,434 | **-4,644** | 0,489 |
| Superior frontal gyrus | 6 | R | 1 | -3 | 72 | -0,286 | 0,64 | 0,81 | **4,576** |
| Precentral gyrus | 6 | L | -39 | -9 | 66 | 1,251 | 1,996 | 0,863 | **4,481** |
| Middle frontal gyrus | 6 | R | 34 | 6 | 45 | **-5,689** | -2,098 | -3,152 | -1,922 |

S1\_3: Affective Task, adults
| Contrasts |  |  |  |  |  | T values |  |  |  |
| --- | --- | --- | --- | --- | --- | --- | --- | --- | --- |
| Region of Interest |  |  | x | y | z | extreme underweight | underweight | normal weight | overweight |
| Cingulate gyrus | 32 | L | -7 | 10 | 34 | 0,618 | 0,356 | -2,286 | 1,477 |
| Insula posterior | 13 | R | 46 | -17 | 18 | **-3,514** | -2,007 | -1,431 | 0,706 |
| Insula anterior | 13 | L | -34 | 5 | 14 | -1,591 | -2,312 | -0,753 | 1,493 |
| Insula posterior | 13 | L | -42 | -15 | 19 | -0,355 | -0,544 | -2,453 | -0,321 |
| Fusiform gyrus | 20 | R | 46 | -29 | -25 | -0,318 | 0,637 | -2,085 | -0,17 |
| Caudate head |  | L | -11 | 11 | 1 | -0,193 | -1,426 | **-4,648** | -1,502 |
| Middle frontal gyrus rostral | 9 | R | 27 | 38 | 25 | 1,353 | -0,325 | 0,143 | -2,159 |
| Middle frontal gyrus | 9 | L | 39 | 38 | 30 | 2,299 | -0,768 | -0,584 | -1,97 |
| Superior frontal gyrus | 10 | L | -34 | 49 | 24 | 1,069 | **-5,365** | **-3,463** | -1,07 |
| Superior frontal gyrus | 10 | R | 32 | 56 | -3 | 0,275 | -0,514 | 0,451 | -2,364 |
| Middle frontal gyrus | 10 | R | 39 | 51 | 10 | 1,21 | 0,001 | -2,501 | **-5,597** |
| Inferior frontal gyrus | 9 | L | -43 | 7 | 30 | -0,516 | -2,981 | **-5,047** | -0,744 |
| Thalamus ventral lateral nucleus |  | R | 12 | -14 | 9 | -1,106 | -0,627 | **-3,376** | 1,627 |
| Medial frontal gyrus | 6 | L | -5 | 2 | 54 | 2,033 | -0,639 | -1,321 | -0,083 |
| Superior frontal gyrus | 8 | L | -18 | 20 | 47 | -0,744 | -0,61 | -2,407 | -1,519 |
| Precentral gyrus | 6 | L | -39 | -9 | 66 | 0,027 | 0,839 | **-3,88** | -1,137 |
| Middle frontal gyrus | 6 | R | 34 | 6 | 45 | **4,633** | 1,47 | 1,012 | -2,044 |

S1\_4: Affective Task, adolescents
| Contrasts |  |  |  |  |  | T values |  |  |  |
| --- | --- | --- | --- | --- | --- | --- | --- | --- | --- |
| Region of Interest |  |  | x | y | z | extreme<br>underweight | underweight | normal<br>weight | overweight |
| Caudate body |  | L | -11 | -20 | 26 | -2,746 | -1,929 | -1,49 | -2,041 |
| Anterior cingulate | 24 | R | 3 | 22 | 10 | 0,731 | **3,817** | 0,974 | -0,403 |
| Cingulate gyrus | 32 | L | -7 | 10 | 34 | -2,746 | -1,929 | -1,49 | -2,041 |
| Insula posterior | 13 | R | 46 | -17 | 18 | **-6,385** | -1,904 | -0,224 | -1,894 |
| Insula anterior | 13 | L | -34 | 5 | 14 | -3,2 | -0,356 | -2,532 | -1,504 |
| Insula anterior | 13 | R | 39 | 6 | 14 | **-4,014** | **-3,376** | -0,24 | -0,721 |
| Insula posterior | 13 | L | -42 | -15 | 19 | **-3,906** | -0,241 | -1,246 | -1,721 |
| Fusiform gyrus | 20 | R | 46 | -29 | -25 | **-3,306** | **-4,615** | -1,432 | 0,018 |
| Fusiform gyrus | 37 | L | -39 | -48 | -9 | **-3,153** | **-3,749** | -0,911 | -1,034 |
| Caudate head |  | L | -11 | 11 | 1 | -1,851 | 1,282 | -2,349 | -0,976 |
| Caudate body |  | R | 14 | -17 | 25 | -2,201 | -2,623 | -0,51 | -3,568 |
| Medial frontal gyrus | 9 | L | -5 | 53 | 38 | -2,472 | -1,584 | -0,668 | -2,098 |
| Inferior frontal gyrus | 9 | R | 50 | 0 | 24 | 1,395 | **3,414** | **4,276** | 0,93 |
| Superior frontal<br>gyrus orbital | 10 | L | -31 | 58 | -2 | **-3,281** | -0,944 | -2,109 | 0,329 |
| Superior frontal<br>gyrus | 10 | R | 32 | 56 | -3 | **-3,221** | -0,941 | -1,552 | 0,34 |
| Medial frontal gyrus | 10 | L | -10 | 65 | 0 | **-3,072** | 0,348 | -0,821 | -0,244 |
| Medial frontal gyrus | 32 | R | 9 | 11 | 45 | **-5,237** | -2,284 | -0,639 | -2,532 |
| Supramarginal gyrus | 40 | R | 61 | -43 | 29 | 3,104 | **3,213** | 1,681 | **5,243** |
| Inferior frontal gyrus | 47 | L | -39 | 25 | 0 | **-5,484** | -2,488 | -1,718 | -0,311 |
| Inferior frontal gyrus<br>rostral | 47 | R | 22 | 35 | -3 | 1,139 | **4,306** | 1,42 | 0,927 |
| Thalamus ventral<br>lateral nucleus |  | R | 12 | -14 | 9 | **-4,694** | 0,04 | -0,641 | 0,385 |
| Medial frontal gyrus | 6 | L | -5 | 2 | 54 | **-4,005** | 0,542 | -0,247 | -0,676 |
| Precentral gyrus<br>medial | 6 | L | -44 | -6 | 41 | **-4,671** | -1,564 | -2,731 | -1,203 |
| Superior frontal<br>gyrus | 8 | L | -18 | 20 | 47 | -1,491 | -0,326 | -2,543 | -0,831 |
| Middle frontal gyrus | 6 | R | 34 | 6 | 45 | -3,51 | -1,049 | -0,638 | -0,068 |

